# Three-dimensional modeling of flow through microvascular beds and surrounding interstitial spaces

**DOI:** 10.1101/2024.02.28.582152

**Authors:** Navaneeth Krishna Rajeeva Pandian, Alanna Farell, Emily Davis, Subramanian Sundaram, Abraham Christoffel Ignatius van Steen, Jessica Li Chang Teo, Jeroen Eyckmans, Christopher S Chen

## Abstract

The health and function of microvascular beds are dramatically impacted by the mechanical forces that they experience due to fluid flow. These fluid flow-generated forces are challenging to measure directly and are typically calculated from experimental flow data. However, current computational fluid dynamics (CFD) models either employ truncated 2D models or overlook the presence of extraluminal flows within the interstitial space between vessels that result from the permeability of the endothelium lining the vessels, which are crucial components affecting flow dynamics. To address this, we present a bottom-up modeling approach that assesses fluid flow in 3D-engineered vessel networks featuring an endothelial lining and interstitial space. Using image processing algorithms to segment 3D confocal image stacks from engineered capillary networks, we reconstructed a 3D computational model of the networks. We incorporated vascular permeability and matrix porosity values to model the contributions of the endothelial lining and interstitial spaces to the flow dynamics in the networks. Simulations suggest that including the endothelial monolayer and the interstitium significantly affects the predicted flow magnitude in the vessels and flow profiles in the interstitium. To demonstrate the importance of these factors, we showed experimentally and computationally that while cytokine (IL-1β) treatment did not affect the network architecture, it significantly increased vessel permeability and resulted in a dramatic decrease in wall shear stresses and flow velocities intraluminally within the networks. In conclusion, this framework offers a robust methodology for studying flow dynamics in 3D in vitro vessel networks, enhancing our understanding of vascular physiology and pathology.

**TRANSLATIONAL IMPACT STATEMENT:** This study introduces a new approach to modeling and flow assessment in 3D microvascular beds and surrounding interstitial spaces. Modeling interstitial space and endothelial monolayer thickness is essential for capturing fluid leakage from the microvascular network into the interstitial space and vice versa when the endothelial monolayer permeability is significantly affected in pathological conditions. Our approach to modeling 3D vascular networks can be used in vivo and in clinical settings to understand flow in tissue microvasculature and its surroundings under disease and healthy conditions.

## Introduction

Continuous blood flow is required throughout the vasculature in the body in order to provide a constant supply of oxygen, nutrients, and immune cells to the tissues. This flow imposes shear and hydrostatic forces on the endothelium lining the vessels, which influences endothelial cell behavior and phenotype [3]. These forces also influence the architecture of the vessel networks, by altering individual vessel diameters and branching, in response to the needs of the tissue for oxygen, nutrients, and trafficking of immune cells [1, 2]. Deviations from the optimal levels of shear and hydrostatic forces can induce pathological changes in cellular phenotype, for example, as observed during atherosclerosis [4, 5].

Despite the importance of these forces to the vasculature, directly measuring them in blood vessels is challenging; instead they are inferred from the flow dynamics and deformations of vessel structures [6]. Animal models are often avoided in these studies due to the complexity, high cost, and the inherent difficulty in measuring flow in vivo. Consequently, in vitro models, such as microfluidic models, are widely employed to study the effects of flow on microvascular networks. Though many microfluidic models are currently available to study the effects of flow on microvascular networks, the exact velocity of flow and forces acting on the micro-vessels of these networks generally are not evaluated [7–9]. Instead, most of these studies only characterize the networks for their architecture (network density, average vessel diameter, average tortuosity of vessels, and more) [8, 10–12]. Recent papers have developed 2D analysis tools to computationally estimate the velocity profiles and corresponding wall shear stresses in such vessel networks, but this approach does not accurately predict flow profiles occurring in complex 3D networks [13, 14]. Some groups have developed 3D computational models of flow, but do not consider the fact that the endothelial cell-based lining of the vascular lumens have a non-zero transmural permeability, allowing flow into the interstitial space between vessels [15, 16]. This interstitial space between vessels in tissues and in 3D culture models is filled with a loose extracellular matrix that provides some resistance to interstitial flow. 2D models have been developed that consider this interstitial space by using a generalized porosity and permeability value for the interstitial space, but have not included the endothelial monolayer itself, which has lower porosity and permeability values than the material of the interstitial space, and therefore should not be neglected [17, 18]. Ultimately, there are currently no 3D flow models that incorporate the interstitial space and/or endothelial lining, despite their potential importance to the resulting flows within these systems.

To overcome these limitations, we developed a bottom-up network modeling approach based on network parameters extracted from 3D image stacks. Our analysis pipeline then estimates the velocities and forces on a 3D in-vitro vessel network by solving the flow equations within the 3-D domain. This model uniquely incorporates both the endothelial monolayer of the vessels and the interstitial space of the tissue as two separate components. Using image processing algorithms in MATLAB, we segment the 3D confocal image stacks obtained from experimental vascular networks in order to extract the network elements’ diameters, lengths, and connectivity. These parameters are then used to construct the network bottom-up, using Boolean operations to reconstruct the network, the endothelial monolayer lining the network, and the interstitial space. This model also incorporates the porosity and permeability values of the endothelial lumen and interstitial space. Finally, we solve the flow equations within the network, the interstitial space, and through the endothelial monolayer. We have validated the model by comparing the predicted velocity values against experimental data.

## Results

### Pipeline to model and analyze microvascular networks

When the dimensions and architecture of the microvascular networks are not predefined, and the microvascular network is formed by self-assembly and morphogenesis in an in vitro system, we need a pipeline to capture and model the network parameters to analyze them. To develop this pipeline, we started by using a microfluidic device with a central tissue chamber flanked by two endothelial cell-lined vessels that enable 3D morphogenesis of an endothelial cell and fibroblast coculture to produce a self-assembled, perfusable vascular network (Figure 1a), similar to a device used previously [8]. Once the networks were formed in the device, they were perfused with fluorescently tagged Dextran, and a stack of images containing different depth positions of the Dextran was created by imaging using a confocal microscope (Figure 1b). We then used several image processing algorithms in Matlab software to convert these image stacks to a 3D model. We used the Dextran image stack and adjusted its aspect ratio to get voxels of equal size in all directions. These adjusted images of the image stack were then converted into binary images. Next, the binarized images were segmented directly by specifying a threshold. These 3D segmentations could be directly exported into volume files that can be used in CFD analysis of flow in the networks but fail to model the interstitial spaces due to the self-intersecting nature of the networks [19–22]. Thus, to generate more network parameters, we used homotopic thinning of the segmentation to generate a 3D medial axis skeleton of the network. This skeleton data was converted into a graph containing links representing the vessel structures and nodes representing the branching and terminal points. The vessel diameters were then calculated based on the distance between the surface and the medial skeleton of the graph links (Figure 1). These parameters were generated at every pixel value, and we took the average values across each link (made up of several pixels) to get smooth surfaces and network elements. If a network link was of high tortuosity (>1.2), we divided the link into five parts to accommodate the tortuosity in the final model.

**Figure 1:**
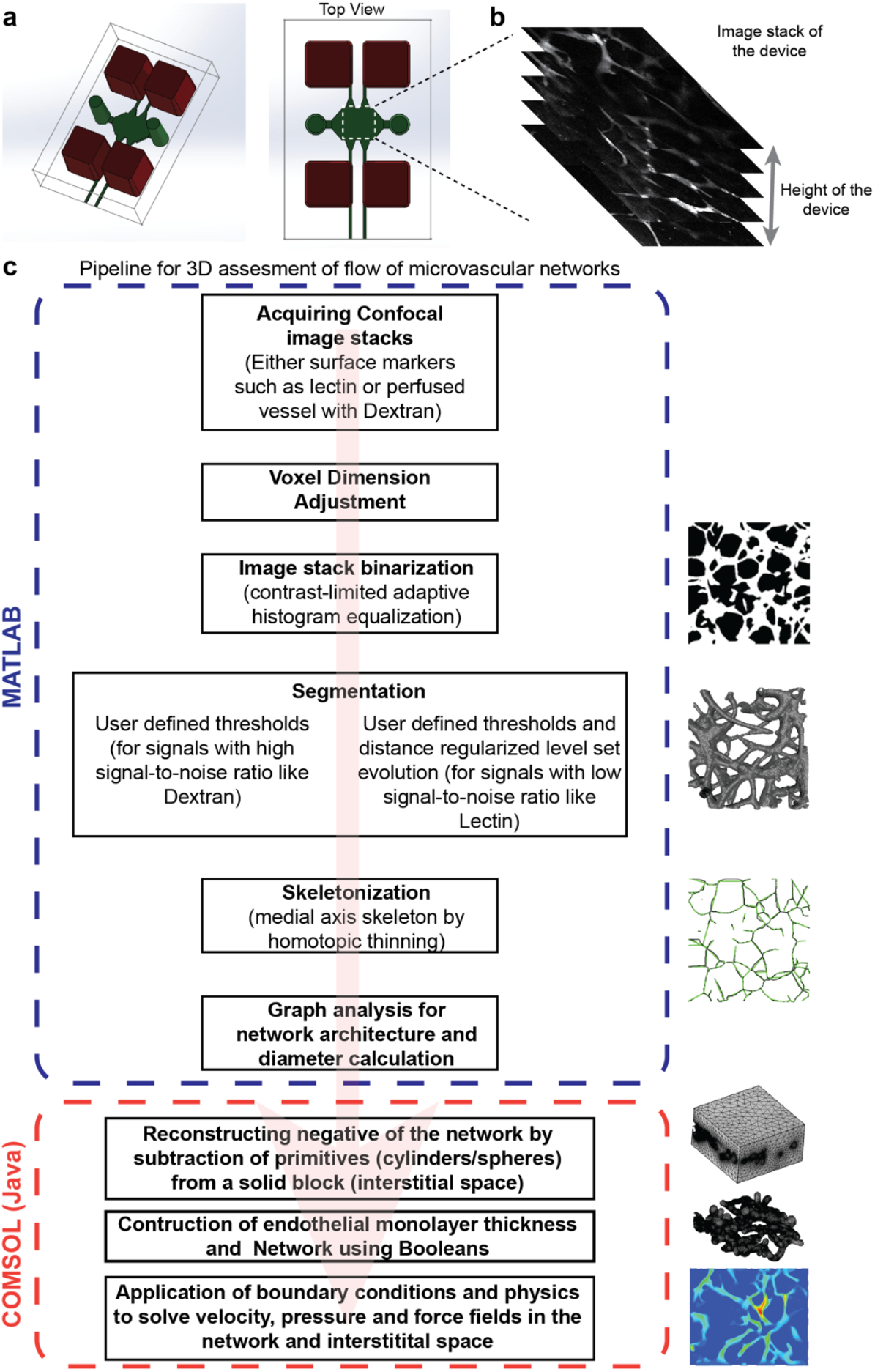
Pipeline to model and analyze microvascular networks. a) Illustration of a two-channel device used to generate self-assembled vascular networks. b) The image stack was prepared from confocal imaging of the networks in the device. c) Pipeline to recreate and analyze the vascular networks using a series of operations in MATLAB (to attain the morphological parameters that define the network’s shape and architecture) and COMSOL (using java for CFD analysis).

After obtaining the vessel morphological parameters, we used a Java (programming language) based algorithm to build the entire network in COMSOL using primitive building blocks such as cylinders, spheres, and blocks. Briefly, we created two negatives of the network in two solid blocks (hollow network in a solid block), which had the dimensions of the interstitial space. One of the negatives is built with the diameters of the network links, and the other has diameters including the endothelial monolayer thickness. The second block is subtracted from the first to obtain a hollow network architecture with endothelial lumen thickness. This hollow network with endothelial lumen thickness is subtracted from a third block (with the exact dimensions as the first and second) to obtain the microvascular network (fluid flow region), endothelial lumen (by retaining the subtracting structure), and interstitial space. In this approach, we place spheres at the ends of the links with corresponding diameters to negate the discontinuities created at the branching points. We then solve the flow equations in free flow and porous media to obtain the flow profiles and forces in the network and its interstitial space. While developing the pipeline, we found that the reconstruction of the interstitial space first instead of the networks reduced errors generated during model building. Once we were able to form a network we visually inspected them and proceeded to validate the model.

### Validation of the models generated by the algorithms

We validated the models generated using the pipeline at two stages. In the first stage, we validated the diameters of the randomly selected individual vessels (links) in the segmentation to that of the raw images. In the second stage, we validated the overall deviation of the architecture of the reconstructed network to the segmentation. We validated the individual link diameter by comparing the computed value to the ground truth manual measurements done on the image cross-sectional projections made in ImageJ (Figure 2a). There was a good agreement between the computed and manually measured values, resulting in a goodness of fit of R^2^ = 0.967. The results of our algorithm are compared with the analysis software REAVER and μVES3D MATLAB environment [16, 23]. Our algorithm has slightly better accuracy than the μVES3D algorithm (with a goodness of fit of R^2^ = 0.857), which uses 3D data for analysis with fewer manual interventions than ours. These 3D-data-based algorithms performed better than the analysis software REAVER (with a goodness of fit of R^2^ = 0.589), which uses 2D data for analysis. The main reason for this is the loss of data in the third dimension when using 2D data sets for measuring the diameters of in vitro microvascular models, which have eccentric vessel cross sections (eccentricity ~0.8) compared to circular in vivo vessels (eccentricity ~1) [16]. We then compared the total volumes of the network models generated by the Matlab algorithm when using different thresholding for binarization with the total volume occupied by pixels above a mean threshold in ImageJ of the raw data. We found that lowering the binarization thresholds gave volumes similar to those predicted from the raw data (Figure 3b). These binarization thresholds for each image varied depending on the overall quality and contrast of the Dextran signal from the surroundings.

**Figure 2:**
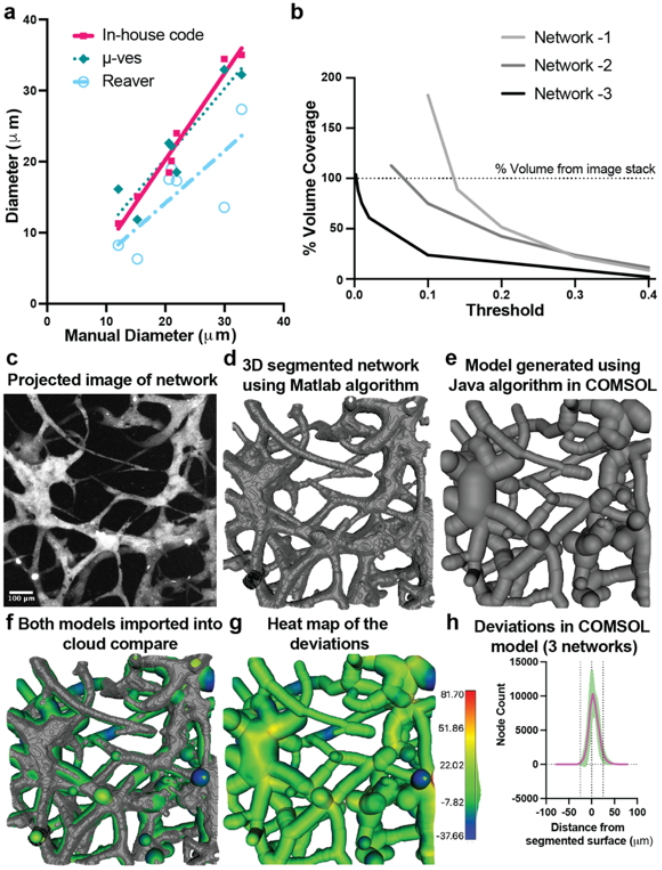
Validation of the models generated by the algorithms. a) Graph comparing the diameters of vessels from the network measured manually with different algorithms (the algorithm used in this work, μ-ves and REAVER). b) Graph showing the convergence of generated volumes with the volume of the raw images for different threshold values. The process of comparing the reconstructed model in Comsol to the raw image is as follows: c) Image of the projection of the network image stack. d) Image of the segmented volume of the network using the Matlab algorithm. e) Image of the volume of the network reconstructed in Comsol using the Java algorithm. f) The image showing the above two volumes imported into Cloudcompare and aligned with each other. g) The heat map showing the deviations on the surface of the volume reconstructed in Comsol compared with the segmented volume using the Matlab algorithm. h) Graph showing the deviations in three reconstructed models.

**Figure 3:**
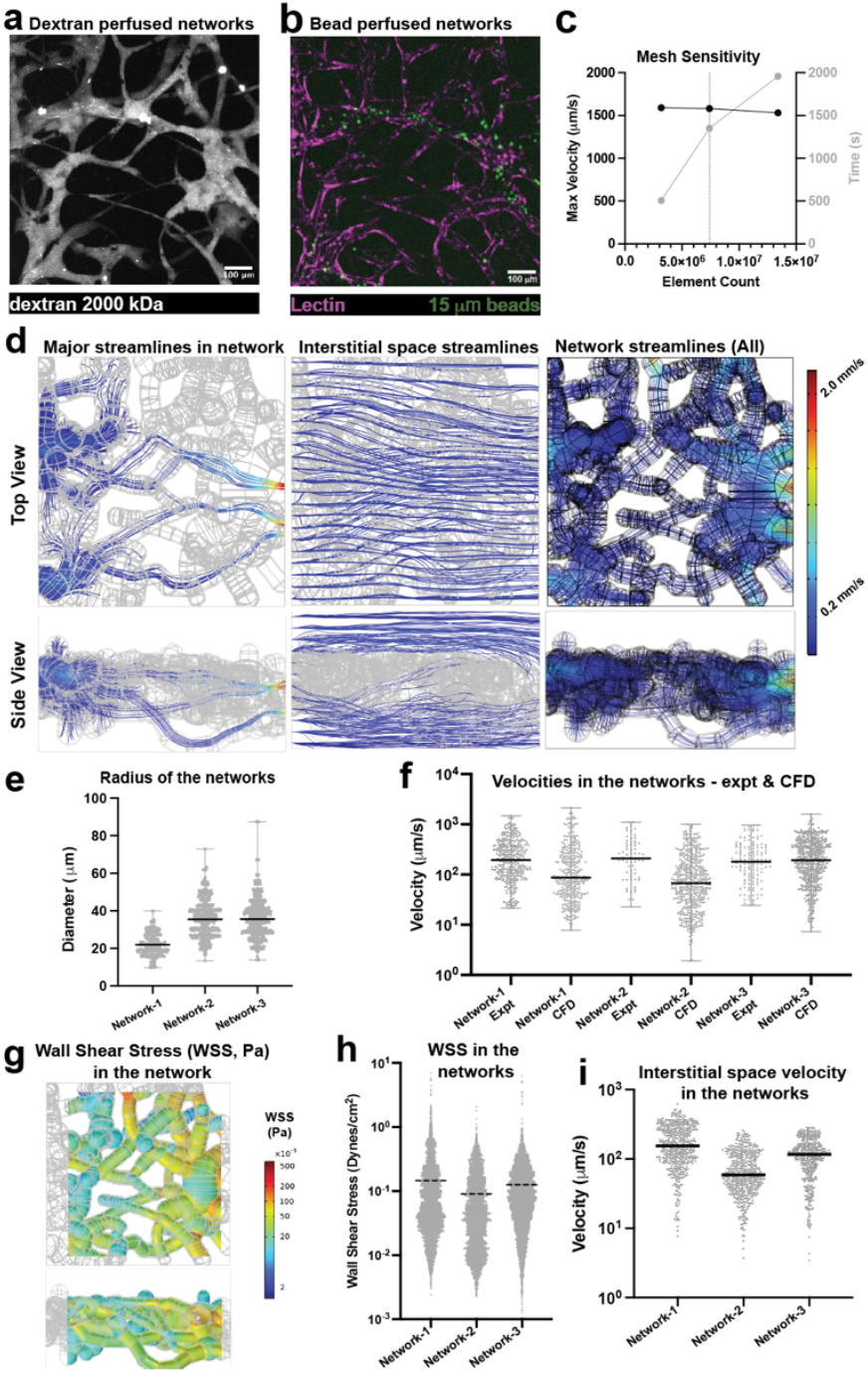
Assessment of in-vitro microvascular networks: a) Image of the network perfused with fluorescently tagged 2000kDa Dextan. b)) Image of the network perfused with fluorescently tagged 10 μm beads and fluorescently labeled lectin. c) graph showing the sensitivity of the model mesh to the maximum velocity predicted in the simulation and the time required for a converged solution. d) Heat maps showing top and side views of the principal streamlines starting from the inlets alone, the whole interstitial space, and the whole network. e) Graph showing radii distribution in the three networks analyzed in this study. f) Graph showing the comparison of the velocities measured in the network from bead perfusion and the CFD analysis for the three networks. g) Heat map showing the wall shear stress in one of the networks. h) Graph showing the wall shear stresses in the network vessels. i) Graph showing the velocity distribution in the interstitial space of the device.

As the Matlab algorithm-generated volumes were similar to the raw data, we next checked if the shape of the networks generated by the Java algorithm correlated with the 3D segmentation (only surfaces are generated) of the networks generated by the Matlab algorithm (Figure 3c & d). The 3D segmented surface from the Matlab algorithm and the network volumes generated from the Java algorithms (Figure 3e) were saved in stereolithography (.stl) format and imported into CloudCompare (a 3D point cloud and mesh processing open-source software, Figure 3f). The models were aligned, and a set of cloud points was generated on the surfaces of each model (as the segmented model is a surface model). The distances between corresponding points on the two models were computed while keeping the points on the 3D segmented surface as the baseline (Figure 3g). We found that 97% of the points in the Comsol model were within 25 μm distance from the 3D segmented surface (Figure 3h). These deviations were mainly caused by the smoothened surfaces created in the bottom-up approach compared to the voxelated surfaces in the segmented model. Also, near the boundaries of the model, the reconstructed model assumed the vessels to have circular cross-sections. In contrast, the segmented model had vessels with noncircular cross-sections by the orthogonal slicing of the vessels at the image boundaries. The smooth surfaces of the final geometry have two advantages, one being that this model is more suitable for computational analysis (avoiding sharp geometric features in the model and, thereby, numerical instabilities due to singularities) and the second being more similar to the actual vascular vessels that do not have sharp features.

### Assessment of in-vitro microvascular networks

Since the reconstructed networks were similar in shape, size, and architecture, we analyzed their flow and compared them with the bead perfusion experimental data. To observe the networks, the devices were perfused with fluorescently tagged large molecular weight Dextran (2000 kDa), which helped maintain the contrast between the networks and interstitial space during imaging (Figure 3a). These networks were then perfused with fluorescently tagged microbeads (~ 10 μm diameter) to measure the flow velocity using Imaris image analysis software (Figure 3b). We used the pipeline to reconstruct a part of the device and discretized the flow and porous domains to solve the flow equations (Figure S1). We then ran a mesh sensitivity analysis to select optimal mesh sizing that gives velocity values within a 5% error with optimal solver time (Figure 3c). We could extract the flow profiles in the microvascular networks and interstitial space (Figure 3d), which showed the 3D distribution of vessels and the corresponding flow profiles. These flow profiles showed that the principal streamlines originating from the inlet vessels followed a path similar to the path the microbeads took in the bead perfusion experiment. We could also map the velocity streamlines in the vessels through which the beads did not perfuse (Figure 3d). These results show that the flow in the interstitial space also has a 3D profile in these in vitro systems.

We analyzed three networks with comparable vessel diameters ranging between 20-40 μm (Figure 3e). We found that the velocities computed by CFD were similar to the velocities measured by the bead perfusion experiment (Figure 3f) and averaged about 100 μm/s for the specified inlet pressure head (5 seconds after having a 4 mm media pressure differential across the device). The average velocity of the networks from the CFD was slightly lower than the corresponding bead perfusion velocity due to the velocities extracted from the vessels in which the beads were not perfused. Once the flow equations were solved to get the flow fields, we could also compute additional parameters, such as wall shear stress and the velocity in the interstitial space of the individual networks (Figure 4g-i). The average wall shear stress in the networks was about 0.1 dynes/cm^2^, which corresponds to the instance at which the bead perfusion was conducted. We can use this knowledge to increase the flow through the system, increasing the shear stress to physiological levels found in the microvasculature (~3-5 dynes/cm^2^) [24]. Thus, we can use the pipeline to compute network parameters such as wall shear stress that cannot be measured directly from an experiment.

**Figure 4:**
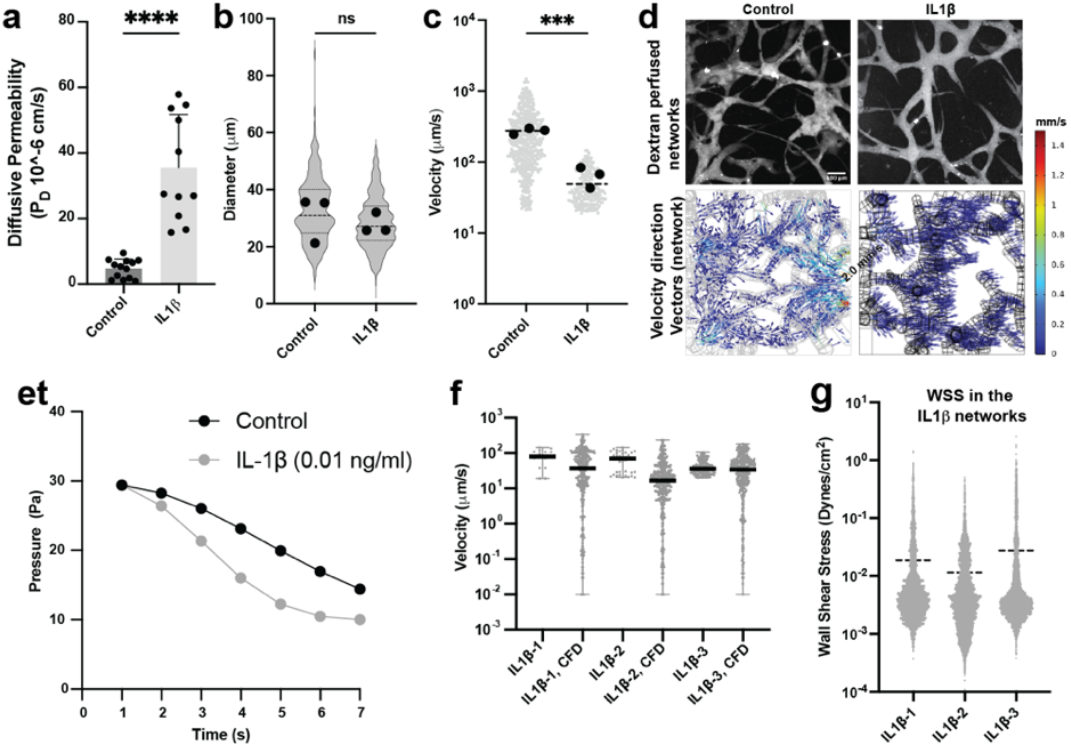
Assessment of in-vitro microvascular networks under cytokine treatment: a) Graph showing the significant difference of 70 kDa Dextran permeability into the fibrin gel in a control and IL-1β treated endothelial lumen in a single channel device (****, p<0.0001). b) Graph showing the differences in the radii distribution in the three control and IL-1β treated networks. c) Graph showing significant difference in the bead perfusion velocities in the three control and IL-1β treated networks (***, p<0.001). d) Graph showing the comparison of the velocities measured in the networks from bead perfusion and the CFD analysis for the three IL-1β treated networks. e) Graph showing the distribution of wall shear stresses in the IL-1β treated network vessels.

### Assessment of in-vitro microvascular networks under cytokine treatment

Next, to assess the performance of our analysis pipeline, we treated our networks with a cytokine to disrupt the endothelial barrier and make it more permeable, thus changing the network and device characteristics. We used IL-1β, a widely known cytokine, to disrupt the endothelial barrier in the network vessels and observe how the flow profiles would be affected. We first treated devices having a single monolayer endothelial lumen (Figure S2, [24]) with IL-1β (0.1 ng/ml for 4 hours), which showed a significant increase in the permeability of the endothelial lumen (Figure 4a). Thus, we treated the network devices with IL-1β for four hours and observed that though there was no significant difference in the morphology of the networks, as evidenced by the diameters of the networks (Figure 4b), we observed a significant difference in the velocity of the beads perfused through them compared to the control untreated networks (Figure 4c). The reduction in velocity is due to two reasons: First, the increased permeability of the endothelial lumen leads to leakage and thus reduced velocity in the vessel (Figure 4d); and second, the increased permeability of the lumen decreases the overall resistance of flow in the networks (Figure 4e), resulting in faster equilibration of media in the media ports at the inlet and outlet, which results in lowering of the flow velocity in the microvessels. We found that the velocities computed by CFD analysis were similar to those measured by the bead perfusion experiment, even though the velocities in these networks had significantly changed (Figure 4f). We also observed reduced wall shear stress in these networks compared to the control untreated networks (Figure 4f compared to Figure 3h). The reduction in the wall shear stress in the networks may have aggravated the loss of barrier function in the microvessels and the effects of the cytokine itself. This experiment shows that inflammatory cytokines (like IL-1β) change flow velocities and wall shear stresses in the microvascular networks and the interstitial spaces, and our analysis pipeline can model these changes accurately.

## Conclusion

This is the first demonstration of the CFD analysis of in-vitro vascular networks that model the 3D microvascular network, its interstitial space, and the endothelial monolayer barrier. Modeling interstitial space and endothelial monolayer thickness is essential as it helps capture fluid leakage from the microvascular network and back into the network from the interstitial space when the endothelial monolayer permeability is significantly affected by cytokines such as IL-1β. We show that changes in endothelial permeability due to cytokines can be incorporated into our system to accurately predict the flow fields and wall shear stresses in the networks. Modeling interstitial space is also crucial as it provides a continuum for the flow between different vessels in the network and automatically solves the in-flow and outflow from network vessels, as illustrated in our study. We also observed that the overall resistance of the device (tissue) decreases for the IL-1β treated condition compared to the control, which shows how a robust endothelial barrier is critical for the proper transport of nutrients and oxygen to the tissues by facilitating enough contact time.

Our modeling approach can be further enhanced with the following improvements. We model the vessels with perfectly rounded cross-sections, whereas the original network vessels have elliptical cross-sections that can be better recapitulated. However, it is worth noting that modeling elliptical cross-sections mayrequire extensive model cleanup for proper discretization. Furthermore, to save modeling time and reduce modeling errors, we have assumed the endothelial monolayer thickness to be uniform all over the network rather than taking the precise thickness of the endothelial lumen at different regions; the spatially varying nature of the thickness can be extracted from the lectin stain images and incorporated into the model if needed.

Overall, this work helps model and analyze 3D vascular networks that can be used in different in vitro and in vivo settings. The 3D model building can also be done from the outputs generated from other network analysis codes such as μVES3D. The model includes microvascular networks, endothelial monolayers, and the tissue’s interstitial space, an essential fluid transport mode in in vitro and in vivo systems. Our approach opens the door to examining the fluid path between blood and lymphatics, where the flow predominantly happens through the interstitial space.

## Methods

### Cell culture

Primary human umbilical vein endothelial cells (HUVEC, CC2519) and normal human lung fibroblasts (HLF, CC-2512). Primary human umbilical vein endothelial cells (HUVEC) from Lonza were cultured in EBM-2 Basal Medium (Cat# CC-3156) with SingleQuots Supplements (Cat# CC-4176). HEK-293T cells were transfected using Lipofectamine 2000 (Invitrogen) with a lentiviral expression vector pLL5.0 and third-generation packaging constructs pMDLg/pRRE, RSV-Rev, and pMD.G. To generate mCherry expressing HUVECs, we transduced HUVECs with puromycin-resistant mCherry in pLL5.0 vectors. The transduced HUVECs were then selected in media containing 2mg/ml of puromycin for two days.

### Single-channel device

The molds for the single-channel microfluidic devices were developed using stereolithography (Proto Labs). Polydimethylsiloxane (PDMS) was mixed at 1:10 ratio and poured on the mold, and cured overnight at 60°C. Individual devices were cut, plasma treated, and bonded to coverslips. To enhance hydrogel bonding to PDMS, the surface inside the device was functionalized with 0.01% poly-L-lysine and 1% glutaraldehyde following plasma activation and washed overnight in DI water. The devices were dried at 100°C for 15 minutes. Acupuncture needles (160 μm diameter) (Hwato) were blocked in 0.1% (w/v) bovine serum albumin (BSA) (Sigma) in phosphate buffer saline (PBS) for 45 minutes and inserted through the two needle guides. Devices with needles were UV-sterilized for 15 minutes. A solution of fibrinogen (2.5 mg/mL), and thrombin (1 U/mL) in EGM-2 was prepared for the bulk hydrogel region of each device. After the addition of thrombin, the solution was quickly injected into the tissue chamber. After 15 minutes, the needles were carefully ablated from the devices to create 300 μm hollow channels between the wells. Each device channel was seeded with additional HUVECs at 2 million cells/mL for at least 5 minutes on each side (top and bottom) in the incubator. Each device received 200 μL of appropriate media daily and cultured on the rocker inside the incubator.

### Network devices

The molds for the 2-channel microfluidic devices were developed using stereolithography (Proto Labs). Polydimethylsiloxane (PDMS) was mixed at 1:10 ratio and poured on the mold, and cured overnight at 60°C. Individual devices were cut, plasma treated, and bonded to coverslips. To enhance hydrogel bonding to PDMS, the surface inside the device was functionalized with 0.01% poly-L-lysine and 1% glutaraldehyde following plasma activation and washed overnight in DI water. The devices were dried at 100°C for 15 minutes. Acupuncture needles (300 μm diameter) (Hwato) were blocked in 0.1% (w/v) bovine serum albumin (BSA) (Sigma) in phosphate buffer saline (PBS) for 45 minutes and inserted through the two needle guides. Devices with needles were UV-sterilized for 15 minutes.

Both HUVECs and Human lung fibroblasts (HLFs, LONZA) were lifted from culture plates using Trypsin (.25%), centrifuged at 200 g for 4 minutes, and resuspended to a concentration of HUVECs (LONZA) - 8.78 million cells/mL in EGM-2 and HLFs – 2.7 million cells/ml. A solution (with final concentrations) of HUVECs (3 million cells/mL), HLFs (1 million), fibrinogen (2.5 mg/mL), and thrombin (1 U/mL) in EGM-2 was prepared for the bulk hydrogel region of each device. After the addition of thrombin, the solution was quickly injected into the tissue chamber, and the devices were repeatedly rotated while the solution was cross-linked. Appropriate media was added to each device well and placed in the incubator (37°C, 5% CO2). After 15 minutes, the needles were carefully ablated from the devices to create 300 μm hollow channels between the wells. Each device channel was seeded with additional HUVECs at 2 million cells/mL for at least 5 minutes on each side (top and bottom) in the incubator. Each device received 200 μL of appropriate media daily and cultured on the rocker inside the incubator.

### Imaging

Devices were fixed using 4% PFA for 15 minutes on a rocker and washed with PBS overnight at 4°C. The devices were then blocked in 3% BSA overnight at 4°C. Lectin (UEA DyLight 649, Vector Labs) was diluted in the PBS at 1:100, added to the devices (for devices not having mCherry (fluorescently) tagged HUVECs), and incubated at 4°C overnight. Devices were washed with PBS and imaged. Just before imaging, 50 ul of 2000 kDa Alexa Fluor-conjugated dextran (0.25 mg/mL) was added to one of the microfluidic channels to generate a pressure gradient between the two channels of the device. All device images were then captured by a Leica SP8 confocal microscope (Leica, Wetzlar, Germany) using a Leica 10x/0.30NA W U-V-I WD-3.60 and Leica LAS X imaging software.

### Image analysis

Images were analyzed or converted and cropped to uniform sizes using Imagej Fiji software before being used in the Matlab software. No other image preparation was done in this manuscript.

### Flow in interstitial space

To measure fluid flow in the interstitial space in the microfluidic devices, Fluorescent Recovery After Photobleaching (FRAP) was used [25]. Devices were incubated with 12.5 ug/ml 70kDa-FITC-dextran (Thermo Fisher Scientific) in PBS for two hours before the experiment. Gel ports in the devices were sealed before the experiment using PDMS (Sylgard 164 1:1). During the experiment, a pressure gradient was generated by pipetting the same 12.5 ug/ml 70kDa-FITC-dextran into the respective media port. Photobleaching of the gel was done on a circle with a radius of 240 μm. Bleaching was performed with the automatic bleaching module (Zeiss Zen software) with the 488 laser at 100%, speed 1, 2 iterations. The device was then imaged every second to track the migration of the bleached area. FRAP experiment was conducted on a confocal LSM980 microscope with Airyscan-2 module (Zeiss) using a Fluar 5x/0.25NA M27 objective (Zeiss).

### Vascular Permeability in single-channel devices

Once a confluent lumen of endothelium was formed in the device, 70 kDa Dextran (Thermo Fisher) tagged with Alexa Fluor 488 was introduced at a concentration of 12.5 *μ*g/ml in the EGM2 medium. This media containing Dextran was introduced into one of the media ports in the device, which was held in a slide holder in a microscope with OKO labs microscope incubator. The diffusion process was imaged every 10 seconds for two minutes with a Zyla 4.2 Plus (sCMOS) camera in a Nikon Ti2-E with Andor Dragonfly (505 System) Spinning Disk Confocal microscope using a 10x 0.4 NA air objective and pinhole CF25. The resulting profile of dextran intensity in the fibrin gel as a function of time was fitted to a dynamic mass-conservation equation as described previously.

The coefficient of diffusive permeability (*P*_D_) was defined by:

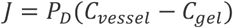

Where *J* is the mass flux of dextran, and *C*_vessel_ and *C*_gel_ are the concentration of Dextran within the vessel and the fibrin gel, respectively. Assuming that the concentration of Dextran is proportional to the intensity of Dextran captured in the images and the initial concentration of Dextran in the gel is negligible, the above equation becomes [24]:

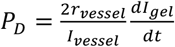

Where *r* is the radius of the vessel (estimated from the image), and *I*_*vessel*_ and *I*_*gel*_ are the intensity of Dextran within the vessel and the fibrin gel, respectively. An existing MATLAB code (Bill’s nature protocol paper) was modified to calculate the diffusive permeability from the timelapse images.

### Bead Perfusion through Vascular Networks

Polystyrene Microspheres, 10 μm, blue fluorescent (365/415) (Fisher Scientific), were added to EGM2 and added to two media ports connecting one of the two parent channels, creating a concentration gradient along networks connecting one parent channel to the other. The device was loaded into the slide holder of a Nikon Ti2-E with Andor Dragonfly (505 System) Spinning Disk Confocal microscope with OKO labs microscope incubator. The bead perfusion process was imaged along the height of the device at every 1 μm height in 0.034 seconds for control networks (0.104 seconds for IL-1ϕ3 treated networks) with a Zyla 4.2 Plus (sCMOS) camera using a 10x 0.4 NA air objective, pinhole CF25 and 100% laser power.

### Bead tracking using Imaris

The image stack of perfusing beads was loaded into ImageJ and converted into an 8-bit image stack for better handling (lowering the file size) of the stack. Beads stuck in the network during the perfusion experiment that may lead to spurious results in the bead tracking process are removed from the image stack by providing appropriate masks. The stack is then saved as an image sequence for further processing in imaris (imaris 10.1). The image sequence tiff files are converted into imaris files using batch processing in the arena (module in Imaris). Next, the imaris file is loaded in the Surpass module, and spots are created for the stack using the spots sub-module. We then use parameters such as spot diameter (10 uM, corresponding to the bead diameter), quality of the spots (20 in this case), mean intensity threshold (depending on the intensity values of a particular image), maximum distance covered (150 um), and number of images to include in a track (three in our case) to filter the spots and their tracks. Once generated, tracks are filtered based on their displacement length (125 um). The track’s mean speeds are exported from specific data selections under detailed statistics for further analysis.

### Comparing Matlab and Java (Comsol) models

Models generated in Matlab and Comsol were exported in stereolithography (.stl) format. These stereolithography models were imported into CloudCompare software. As the model in Matlab was generated from the image file before scaling correction, it had dimensions in pixel units 255x255xh μm. The model in Comsol was 906x906x(h*slice height) μm. The Matlab model was translated and scaled in CloudCompare using the scaling parameters to have the same centroid as the Comsol model. The two models are registered using the fine registration tool by making the Matlab model the reference and root mean square value (RMS value) for comparison as 1e-5. Then, using the cloud-to-mesh tool we compare the overlap between the two models.

### Computational model

We developed the 3D steady-state model using the CFD module in the Comsol software. The computational domain comprised the fibrin gel, lumen, and the lumenized network. We solved for the conservation of mass and Navier Stoke’s equations in the vessel networks where bulk fluid flow occurs. Assuming the fluid to be incompressible:

Conservation of mass

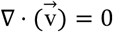

Navier Stoke’s equation.

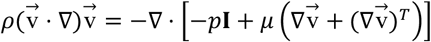

Where *ρ* is the fluid density (1000 kg m^−3^ for media), *μ* is the fluid viscosity (1.002 × 10^−3^ Pa s). 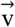 is the velocity vector, and *p* is the pressure (scalar) in the networks. The fibrin interstitial space (also containing fibroblasts) and the endothelial lumen were modeled as porous media, and we solved for the Brinkman equations with Forchheimer correction and conservation of mass to describe the flow:

Brinkman equations with Forchheimer correction.

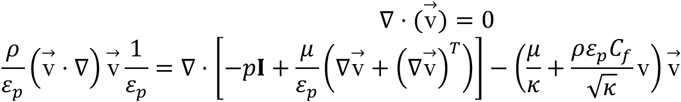

Where *κ* (m^2^) is the permeability, and *ε*_*P*_ is the porosity of the interstitial space or endothelial lumen. The dimensionless friction factor in the porous is related to its porosity (*ε*_*P*_).

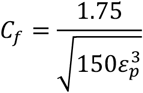

As the above equations reveal, the momentum transport equation in the free and porous media are closely related. The Brinkman equation replaces the convection-dependent momentum transfer in free flow by the drag force experienced by the fluid in the porous media. The last term in the Brinkman equation is the Forchheimer correction for turbulent drag contributions.

In this study, we have assumed *ε* _*P*_ to be 0.265 for both the fibrin hydrogel and the endothelial monolayer thickness. Also, *κ* (m^2^) is assumed to be 1.2e-7 [26] for the fibrin gel region, and 5.73e-13 for the control endothelial monolayer, and 5.73e-12 for IL-1β treated devices.

In this study, we solved the steady-state flow equations for pressure inlet conditions. The pressure at the inlet of the vessels and the fibrin interstitial space of the control were assumed to be 3.15 Pa and 1.75 Pa, respectively. The pressure at the inlet of the vessels and the fibrin interstitial space of the control were assumed to be 0.35 Pa and 0.075 Pa, respectively. The outlet boundary conditions are assumed to be 0 Pa pressure boundary conditions. The inlet pressure values were chosen from the CFD analysis of a chip with two parent channels of 250 μm diameter and a network of 50 μm diameter vessels, making a predefined square grid (Figure S3).

### Statistical Testing

For all experiments, independent two-sample populations will be compared using unpaired, non-parametric two-sample t-tests. \**P* < 0.05 was considered to be statistically significant. Statistical analyses were performed using GraphPad Prism software.

## Supporting information

Supplement Images

## Funding and acknowledgments

This work was supported by the NIH (EB033821), NSF STC CEMB, BSF (2017239), and the Wellcome Leap HOPE Program C. S. C.

## Ethics Statement

All research conducted in this study adhered to the ethical principles outlined in the Declaration of Helsinki.

