## Supplement Images for "Three-dimensional modeling of flow through microvascular beds and surrounding interstitial spaces"

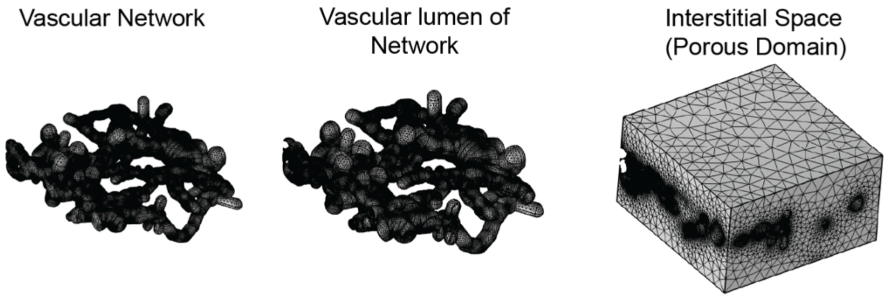


*Figure S 1: Discretized vascular network, vascular lumen of the network, and the interstitial space modeled as a porous domain.*

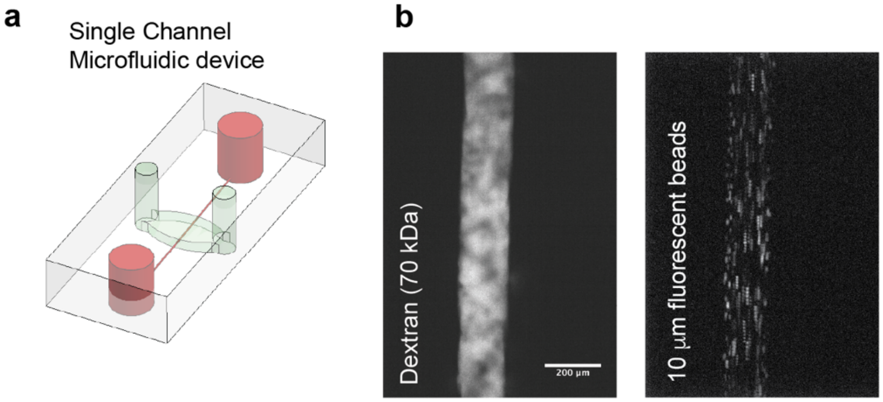


*Figure S 2: a) A single channel device is used to measure the diffusive permeability of the vascular lumen. b) Dextran (70 kDa) and fluorescent microbeads perfused through the single channel device.*

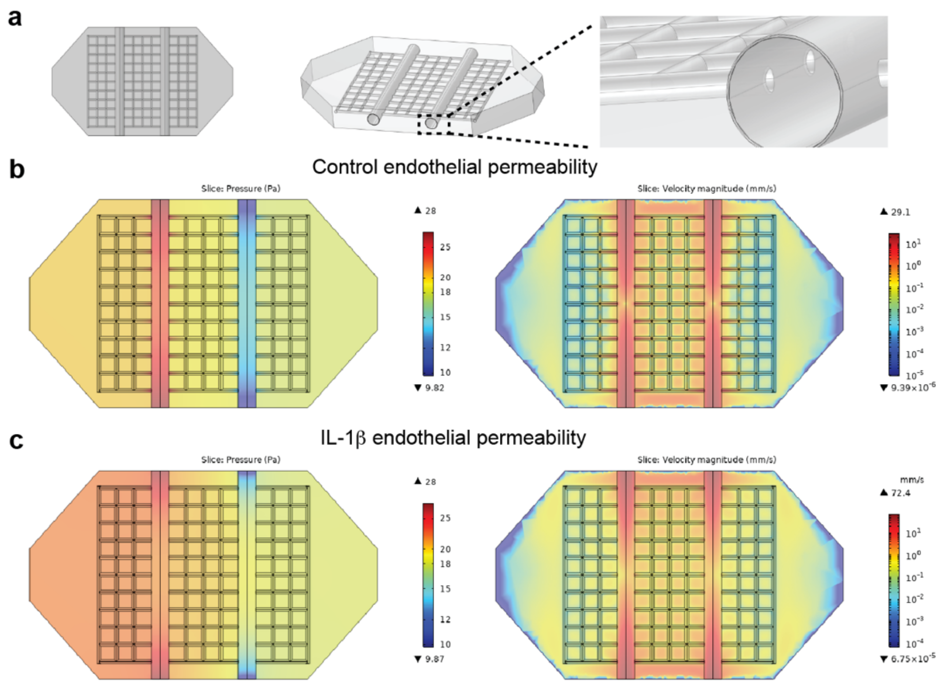


*Figure S 3: A square grid network model used for calculating the inlet pressure conditions a) model of the square grid matrix b) pressure and velocity heat maps of control endothelium for the grid network c) pressure and velocity heat maps of IL1β treated endothelium for the grid network.*
